# A pre-vertebrate endodermal origin of calcitonin-producing neuroendocrine cells

**DOI:** 10.1101/2024.02.17.580846

**Authors:** Jenaid M. Rees, Katie Kirk, Giacomo Gattoni, Dorit Hockman, Dylan J. Ritter, Èlia Benito-Gutierrez, Ela W. Knapik, J. Gage Crump, Peter Fabian, J. Andrew Gillis

**Author notes:** Equal contribution (Jenaid M. Rees, Katie Kirk, Giacomo Gattoni, J. Andrew Gillis).

## Abstract

Vertebrate calcitonin-producing cells (C-cells) are neuroendocrine cells that secrete the small peptide hormone calcitonin in response to elevated blood calcium levels. C-cells are crucial for maintenance of calcium homeostasis, yet the embryonic and evolutionary origins of this cell type remain contentious. Whereas mouse C-cells reside within the thyroid gland and derive from pharyngeal endoderm, avian C-cells are located within ultimobranchial glands and were reported to derive from the neural crest. We use a comparative cell lineage tracing approach in a range of vertebrate model systems to resolve the ancestral embryonic origin of vertebrate C-cells. We find, contrary to previous studies, that chick C-cells derive from pharyngeal endoderm, with neural crest-derived cells instead contributing to connective tissue intimately associated with C-cells in the ultimobranchial gland. This endodermal origin of C-cells is conserved in a ray-finned bony fish (zebrafish) and a cartilaginous fish (the little skate, *Leucoraja erinacea*). Furthermore, we discover putative C-cell homologues within the endodermally-derived pharyngeal epithelium of the ascidian *Ciona intestinalis* and the amphioxus *Branchiostoma lanceolatum*, two invertebrate chordates that lack neural crest cells. Our findings point to a conserved endodermal origin of C-cells across vertebrates and to a pre-vertebrate origin of this cell type along the chordate stem.

## Introduction

Calcitonin-producing neuroendocrine cells (“C-cells”) are a specialized cell type of vertebrate animals that secretes the small peptide hormone calcitonin in response to elevated blood calcium (hypercalcaemia)^1–4^. In bony vertebrates, macrophage-like cells called osteoclasts resorb and remodel bone, releasing calcium into the bloodstream. Calcitonin from C-cells, in turn, lowers blood calcium levels by inhibiting osteoclast activity^5,6^ and promoting calcium deposition within bone^7^. Calcitonin is widely used for the acute treatment of metabolic bone disorders, such as osteoporosis^8^ and Paget’s disease^9^ – though, paradoxically, humans with varying levels of endogenous calcitonin (e.g., persons that have undergone thyroidectomy or with C-cell-derived medullary thyroid carcinoma/MTC) exhibit no differences in bone mineral density^10,11^. Nevertheless, osteoclast function is highly sensitive to calcitonin levels *in vitro*^12^, and a mouse model of calcitonin deficiency (i.e., the *Calca*^-/-^ mouse, which lacks the gene encoding both calcitonin and its splice variant, calcitonin gene-related peptide) exhibits increased bone resorption with aging and decreased bone mineral density during lactation^13,14^. In cartilaginous fishes (sharks, skates and rays), calcitonin is reported to decrease blood calcium levels in some taxa^15^, and increase blood calcium in others^16^. Cartilaginous fishes lack bone and osteoclasts, but in these fishes, calcitonin may act on the gallbladder to regulate calcium concentrations and excretion in bile^17^. Calcitonin therefore seems to function as a conserved modulator of calcium homeostasis in vertebrates – and possibly more broadly among animals^18^ – though its mechanism(s) of action and responsive cell types remain incompletely understood outside of mammals^19^.

In jawed vertebrates, C-cells develop within the ultimobranchial bodies (UBs): paired structures that derive from caudal pharyngeal pouches during early craniofacial development^20^. Mammalian UBs are transient structures that develop from the fourth pharyngeal pouches and that ultimately migrate toward the midline to merge with the developing thyroid primordium^21^. This accounts for the final location of mammalian C-cells (also known as parafollicular cells) within the thyroid gland. In non-mammalian vertebrates (i.e., birds, reptiles, amphibians, and fishes), UBs do not fuse with the thyroid gland, and instead persist as distinct, paired C-cell-containing glands in the neck or caudal pharynx^22–24^.

Though the location of C-cells within the UB (or mammalian thyroid gland) is well established, the embryonic origin of the cell type has been a matter of debate for over 50 years. Pearse, Polak, LeDouarin and colleagues reported a neural crest origin of avian C-cells based on a series of quail-chick chimaera lineage tracing experiments^25–27^. Following isotopic transplantation of quail neural tube into chick host embryos, they observed quail neural crest-derived cells within the host ultimobranchial gland that appeared to exhibit cytoplasmic secretory granules characteristic of C-cells^25^, formaldehyde-induced fluorescence characteristic of some neuroendocrine cell types^26^, and positive immunostaining with an anti-calcitonin antibody^27^. These findings led to assumption of a neural crest origin of C-cells in all vertebrates (including mammals), of C-cell-derived cancers like MTC^28–30^, and more broadly, of other biochemically similar neuroendocrine cell types (the amine precursor uptake and decarboxylation or APUD series) that are distributed throughout the body in disparate organs and tissues^31–33^.

But in the decades following reports of a neural crest origin of avian C-cells, corroborating evidence for a neural crest origin of mammalian C-cells remained scant. Kameda and colleagues tested for a neural crest contribution to C-cells in mouse by Conexin43-lacZ or Wnt1-Cre lineage tracing and found none – rather, they reported that developing mouse C-cells express E-cadherin (a common marker of epithelial cell types)^34^. More recently, Johanson *et al.* demonstrated unequivocally by Sox17-Cre lineage tracing that mouse C-cells derive from pharyngeal endoderm, and not from the neural crest^35^. The distinct germ layer origins of C-cells in mouse and chick could indicate that these cell types are not homologous (i.e., that mammals and birds independently evolved calcitonin-secreting neuroendocrine cells in their thyroid glands and UBs, respectively), or that C-cells have undergone a radical lineage shift – from endoderm to neural crest, or vice versa – during tetrapod evolution.

While it is now recognized that many other neuroendocrine cell types within Pearse’s APUD series have non-neural crest embryonic origins (e.g., pancreatic endocrine cells^36^, pulmonary neuroendocrine cells^37^ and gut enterochromaffin cells^38^), a conserved neural crest origin of C-cells remains cemented in the medical literature, despite a dearth of data on the embryonic origin of C-cells from other taxa. Here, we resolve the ancestral embryonic origin of C-cells using *in situ* gene expression analysis and cell lineage tracing in a range of vertebrate model systems. We revisit the embryonic origin of avian C-cells in the chick and find that while neural crest cells do contribute connective tissue to the UBs, they do no give rise to C-cells. Rather, chick C-cells derive from endoderm, as in the mouse. We also find that an endodermal origin of C-cells is conserved in a ray-finned bony fish (zebrafish) and in a cartilaginous fish (the skate, *Leucoraja erinacea*). Finally, we test for calcitonin expression in two invertebrate chordate taxa that lack neural crest cells – the ascidian *Ciona intestinalis* and the amphioxus *Branchiostoma lanceolatum* – and we identify putative C-cell homologues within their endodermally-derived pharyngeal linings. These findings point to an ancestral and conserved endodermal origin of C-cells within chordates and broaden the repertoire of endodermal cell types in the ancestral chordate.

## Results and Discussion

### Chick C-cells derive from endoderm, and not neural crest

Considering recent genetic lineage tracing data showed an endodermal origin of C-cells in mouse, we decided to revisit the germ layer origin of avian C-cells. In embryonic day (E)10 chick embryos, the UBs are located between the esophagus and the carotid artery, at the level of the nodose ganglion (Fig. 1a), and chick C-cells express both calcitonin (Fig. 1b) and tyrosine hydroxylase (Fig. 1c; Fig. S1). The neural crest origin of chick C-cells was previously reported based on cell lineage tracing using chick-quail chimeras^25–27^. We carried out similar experiments, labeling premigratory cranial and vagal neural crest cells by unilaterally grafting cranial and vagal-level neural fold from GFP+ transgenic donor chick embryos^39^ to wild-type hosts. Grafted embryos were grown to embryonic day (E)8.5-10, and then sectioned and immunostained for GFP and calcitonin. We analysed three embryos that received grafts of GFP+ neural fold from the otic vesicle to somite 1, and three embryos that received grafts of GFP+ neural fold from the otic vesicle to somite 7 (Fig. 1d). In all six embryos, we recovered GFP+ cells inside and around the UB but no colocalization of GFP and calcitonin in any embryos (Fig. 1e; Fig. S2). This strongly suggests that chick C-cells do not derive from the neural crest, and that previous lineage tracing experiments using quail-chick chimeras may have recovered a neural crest contribution to connective tissue within the UB, rather than to the calcitonin-expressing C-cells.

**Figure 1:**
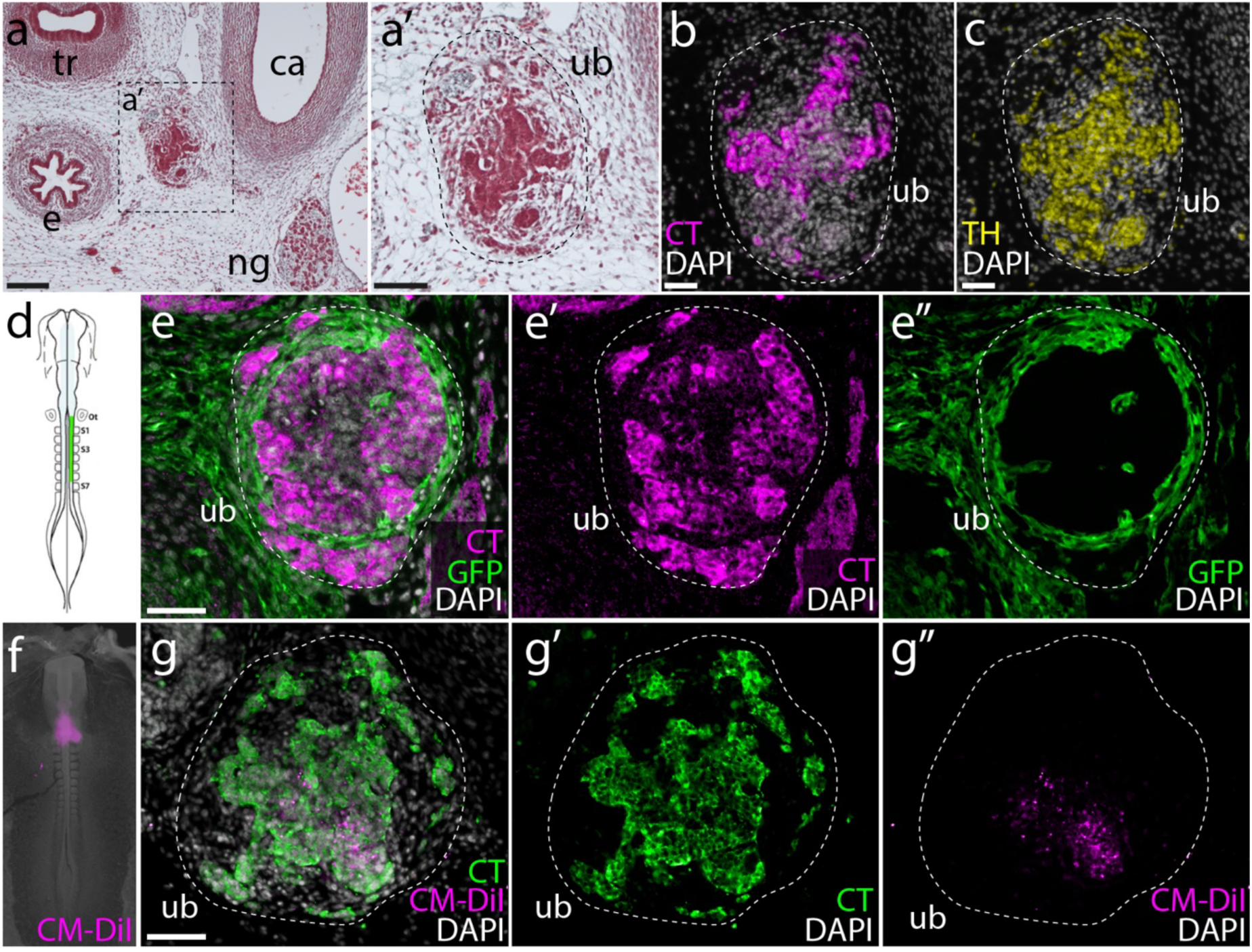
Chick C-cells derive from endoderm, not neural crest. **a,a’)** At E10, the chick UB is located between the esophagus and the carotid artery, at the level of the nodose ganglion. Chick C-cells co-express **b)** calcitonin (CT) and **c)** tyrosine hydroxylase (TH). **d)** Neural crest lineage tracing was performed by isotopic unilateral grafting of GFP+ neural fold into a wild-type host embryo. In this example, a graft was performed with neural fold from the otic vesicle to S7. **e)** In grafted embryos, we recovered GFP+ cells in and around the UB, but we observed no co-localisation of GFP and CT. **f)** Endodermal lineage tracing was performed by microinjecting CM-DiI into the pharyngeal cavity of chick embryos at HH11. **g)** In three out of five labeled embryos, we recovered CM-DiI within the UB at E8.5, with co-localization of CM-DiI and CT indicating endodermal origin of chick C-cells. *ca*, carotid artery; *e*, esophagus; *ng*, nodose ganglion; *ot*, otic vesicle; *S1-7*, somites 1-7; *tr*, trachea; *ub*, ultimobranchial body. Scale bars: **a** = 100 μm; **a’** = 50 μm; **b,c,e,g** = 25 μm.

To test for an alternative endodermal origin of C-cells in the chick, we microinjected the lipophilic dye CM-DiI into the pharyngeal cavity of chick embryos at Hamburger Hamilton stage 11 (Fig. 1f), to label the endodermal epithelium that lines the pharyngeal cavity. We then grew injected embryos to E8.5 and tested for co-localisation of CM-DiI and calcitonin by immunofluorescence. In 3/5 labeled embryos, we recovered CM-DiI-labeled C-cells within the UBs (Fig. 1g), indicating an origin of these cells from pharyngeal endoderm. So, while the UB is surrounded by and contains some neural crest-derived cells, we find no evidence that C-cells themselves derive from the neural crest in chick. Instead, as in mouse, we find that chick C-cells have an endodermal origin.

### Zebrafish C-cells derive from endoderm

We next sought to resolve the embryonic origin of C-cells in a ray-finned fish outgroup to the tetrapods. This would allow us to infer the ancestral germ layer origin of C-cells for bony vertebrates. In the larval zebrafish, the UBs meet at the ventral midline beneath the ventral wall of the esophagus, at the axial level of the sinus venosus (Fig. 2a). C-cells within the UB are recognizable by their expression of *calca* (with splice variants of *calca* encoding calcitonin and calcitonin gene-related peptide) by mRNA *in situ* hybridization (Fig. 2b). We analyzed *tfap2a^mob^;foxd3^mos^* zebrafish, which lack neural crest derivatives^40^, to determine whether these mutants also lack C-cells. We first tested for *calca* expression in cells of the developing UB at 7 days post fertilization (dpf) in wild-type siblings, and we observed *calca*+ cells within the UB of all larvae (Fig. 2c,d; *calca*+ cells found in n=6/6 individuals analyzed). At 7 dpf, *tfap2a^mob^;foxd3^mos^* mutants exhibit a distinct lack of pigment and craniofacial malformations (Fig. 2e), owing to an absence of neural crest-derived melanocytes and skeletal tissues, respectively, but we found that these mutants nevertheless possessed *calca*+ cells within their UB (Fig. 2f; *calca*+ cells found in n=6/6 individuals analyzed). The presence of *calca*+ cells in the UBs of mutants lacking neural crest cells indicates a likely non-neural crest origin of C-cells in zebrafish.

**Figure 2:**
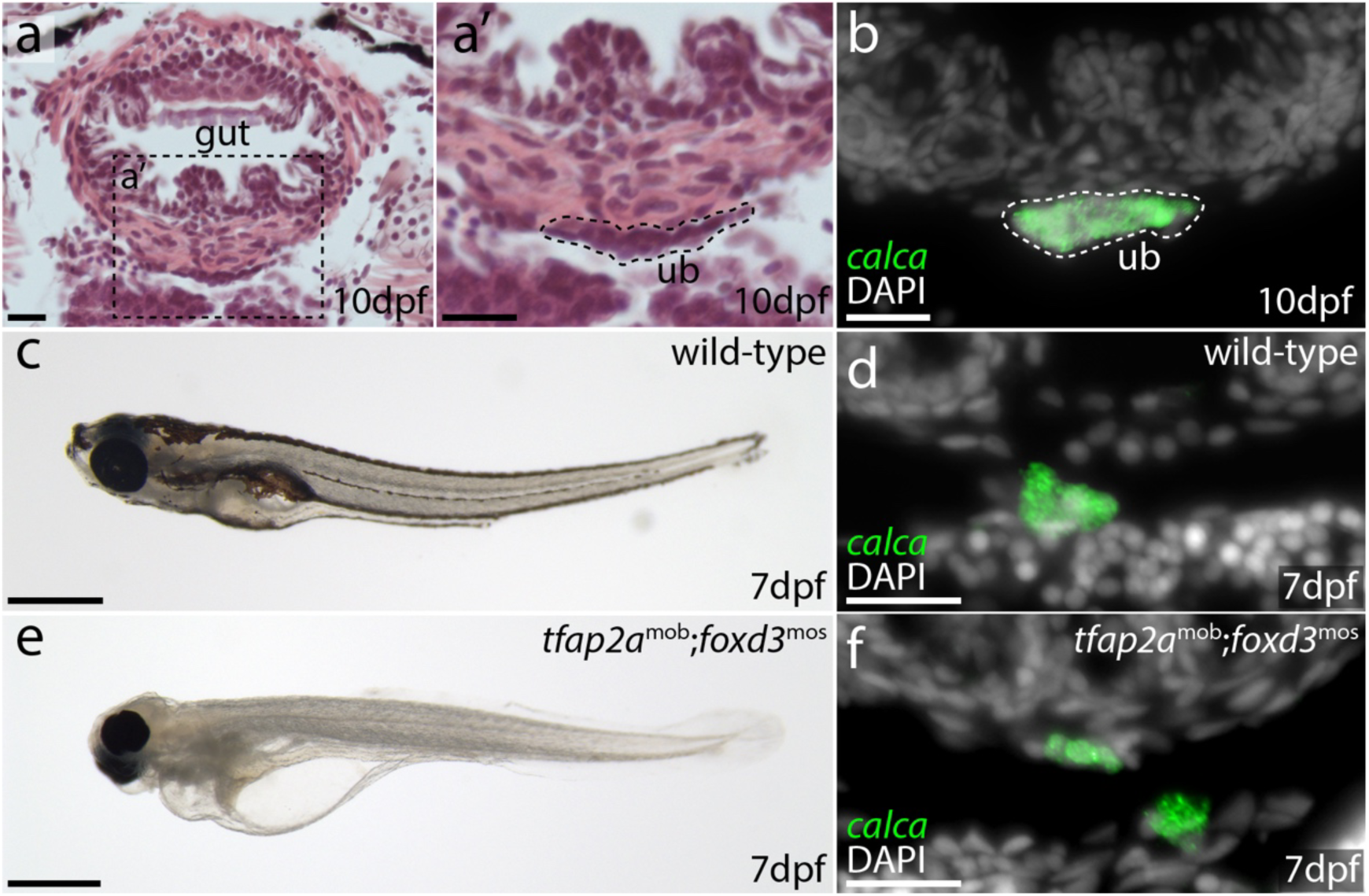
C-cells of the larval zebrafish develop in the absence of neural crest. **a)** The UB of a 10 dpf zebrafish is located between the gut and the heart, at the axial level of the sinus venosus. It is recognizable as **a’)** a distinct cluster of cells beneath the muscle layer of the gut, and **b)** by its expression of *calca*. **c)** 7 dpf wild type zebrafish possess **d)** *calca*+ C-cells within their UB. **e)** 7 dpf *tfap2a*^mob^;*foxd3*^mos^ mutants lack neural crest cells, but **f)** still possess *calca*+ C-cells in their UB. *ub*, ultimobranchial body. Scale bars: **a,a’,b,d,f** = 20 μm; **c,e** = 1 mm.

Next, we employed genetic cell lineage tracing to test for a neural crest contribution to C-cells in the ultimobranchial gland of zebrafish. Although we were able to detect transcription of *calca* in the UB of larval zebrafish from 7-10dpf, we were unable to detect expression of calcitonin protein at these stages. We therefore opted to test for neural crest contributions to UB C-cells in adult zebrafish when calcitonin expression is readily detectable by immunofluorescence. We used Sox10:Cre; actab2:loxP-BFP-STOP-loxP-dsRed (Sox10>dsRed) fish, which enabled the permanent labeling of neural crest cell derivatives shortly after their differentiation at 10 hours post-fertilization^41,42^. Fish were raised to adulthood (150-210 dpf), and sections through the entire UB were co-stained for the dsRed reporter of neural crest lineage and calcitonin by immunofluorescence. In 3/3 fish examined, we observed no dsRed+ cells within the UBs (Fig. 3a), despite strong labeling of other neural crest derivatives within the same sections (e.g. TH+ neurons of the sympathetic ganglia – see inset boxes in Fig. 3a). These observations further support a non-neural crest origin of C-cells in zebrafish.

**Figure 3:**
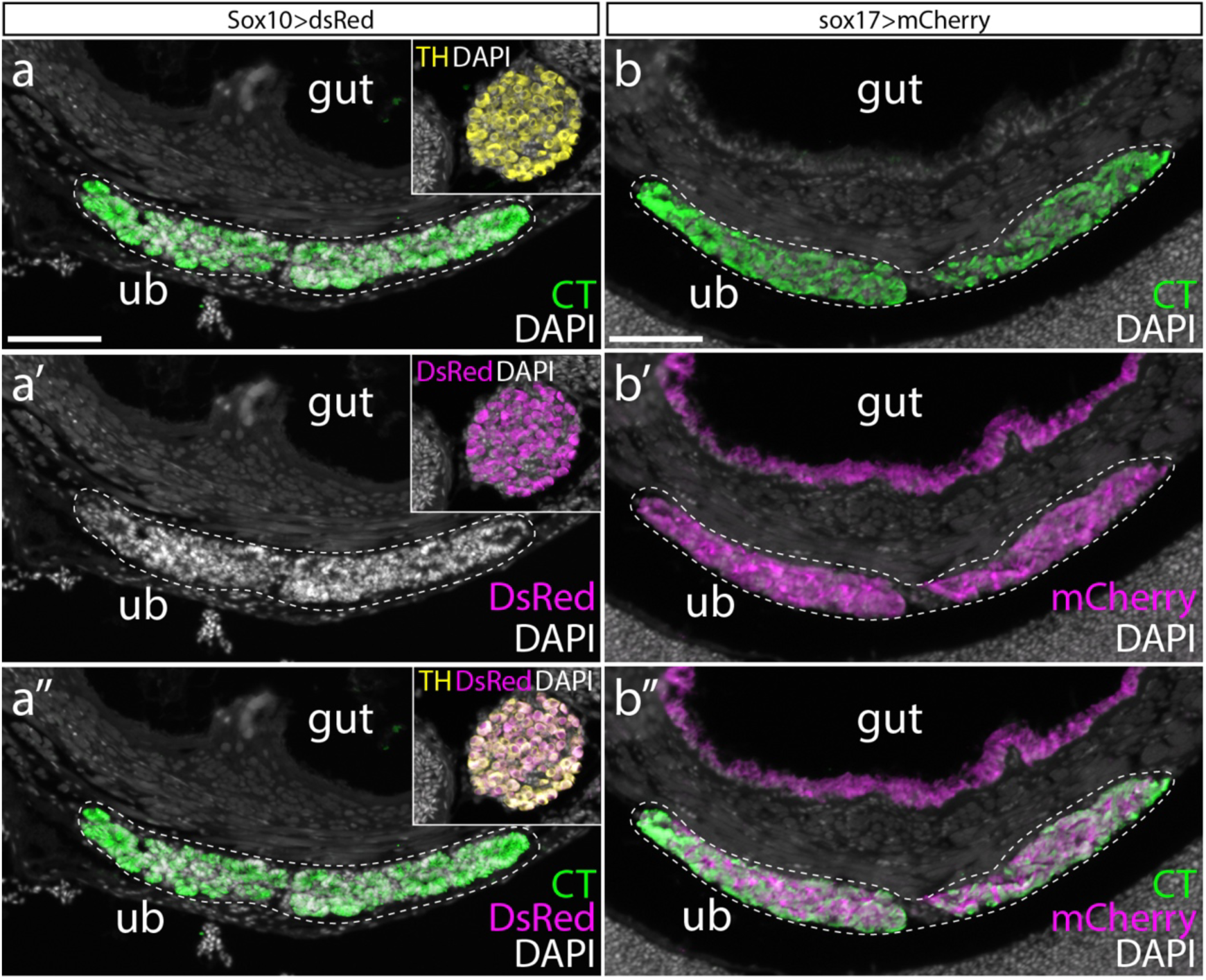
Zebrafish C-cells derive from pharyngeal endoderm, and not neural crest. **a)** The UB of adult zebrafish stains positively for calcitonin (CT) by immunofluorescence. The Sox10>dsRed line labels all neural crest derivatives with DsRed. We found no DsRed+ cells within the UB of adult fish of this line, despite strong labeling of other neural crest-derived strictures on the same sections – e.g. Tyrosine Hydroxylase (TH)-positive neurons of the sympatheic ganglia (see inset boxes in **a**, **a’** and **a”**). **b)** sox17>mCherry zebrafish embryos were treated with 4OHT during gastrulation to indelibly labeling the endodermal lineage with mCherry, and then grown to adult. We observed complete colocalization of mCherry and CT within the UB of adult fish, indicating endodermal origin of zebrafish UB C-cells. *ub*, ultimobranchial body. Scale bars: **a,b** = 50 μm.

Having excluded neural crest cells as progenitors for C-cells, we tested for an endodermal origin of zebrafish C-cells. To specifically mark endodermal lineages in zebrafish, we employed sox17:CreERT2;ubi:loxP-eGFP-STOP-loxP-mCherry (sox17>mCherry) fish and treated them with 4-hydroxytamoxifen (4OHT) during gastrulation (4-6 hours post-fertilization) to induce loxP recombination and indelible labeling of the endodermal lineage with mCherry^43^. Fish were raised to adulthood (150-210 dpf), and sections through the UBs were co-stained for the mCherry reporter of endodermal lineage and Calcitonin by immunofluorescence. We found consistent co-expression of the mCherry reporter of endodermal lineage and calcitonin throughout the UBs in 3/3 fish analyzed (Fig. 3b), indicating an endodermal origin of C-cells in zebrafish. When considered alongside data from mouse and chick, this finding indicates that C-cells ancestrally derive from endoderm in bony vertebrates.

### Skate C-cells derive from endoderm

All extant jawed vertebrates belong to one of two lineages: bony vertebrates (including bony fishes and tetrapods) and cartilaginous fishes (sharks, skates, rays and holocephalans). To test whether an endodermal origin of C-cells is conserved in cartilaginous fishes, we carried out a series of cell lineage tracing experiments in a cartilaginous fish outgroup to the bony vertebrates, the little skate (*Leucoraja erinacea*)^44^. In skate embryos, UBs are located between the caudal gill arches and the pectoral girdle (Fig. 4a), and as in bony vertebrates, skate C-cells within the UBs are marked by expression of calcitonin (Fig. 4b). To test whether skate C-cells derive from endoderm, we microinjected the pharyngeal cavity of neurula-stage skate embryos with the lipophilic dye CM-DiI (Fig. 4c). By microinjecting CM-DiI into the pharyngeal cavity before pharyngeal pouches and gill slits have formed, we can broadly label pharyngeal endoderm without contaminating adjacent tissues (Fig. 4d)^45,46^. Injected embryos were then left to develop until stage (S)32 (∼8-10 weeks post-injection), by which point the UBs are differentiating. Embryos that received pharyngeal endodermal labeling with CM-DiI showed abundant CM-DiI-retention throughout the developing UBs (Fig. 4e). To specifically test for an endodermal origin of C-cells, we sectioned twelve S32 embryos that had received CM-DiI-labelling of pharyngeal endoderm and found that eight embryos showed co-localization of CM-DiI with *Calca* expression within the UB (Fig. 4f-g). We also examined 20 skate embryos that received microinjection of CM-DiI into the neural tube at neurula stage (to label pre-migratory neural crest cells), and we found no CM-DiI-labeled cells in the UB of any of these embryos^47^. These findings indicate that C-cells derive from pharyngeal endoderm in the skate. When considered alongside data from mouse and zebrafish, this finding supports an ancestral endodermal origin of C-cells for jawed vertebrates.

**Figure 4:**
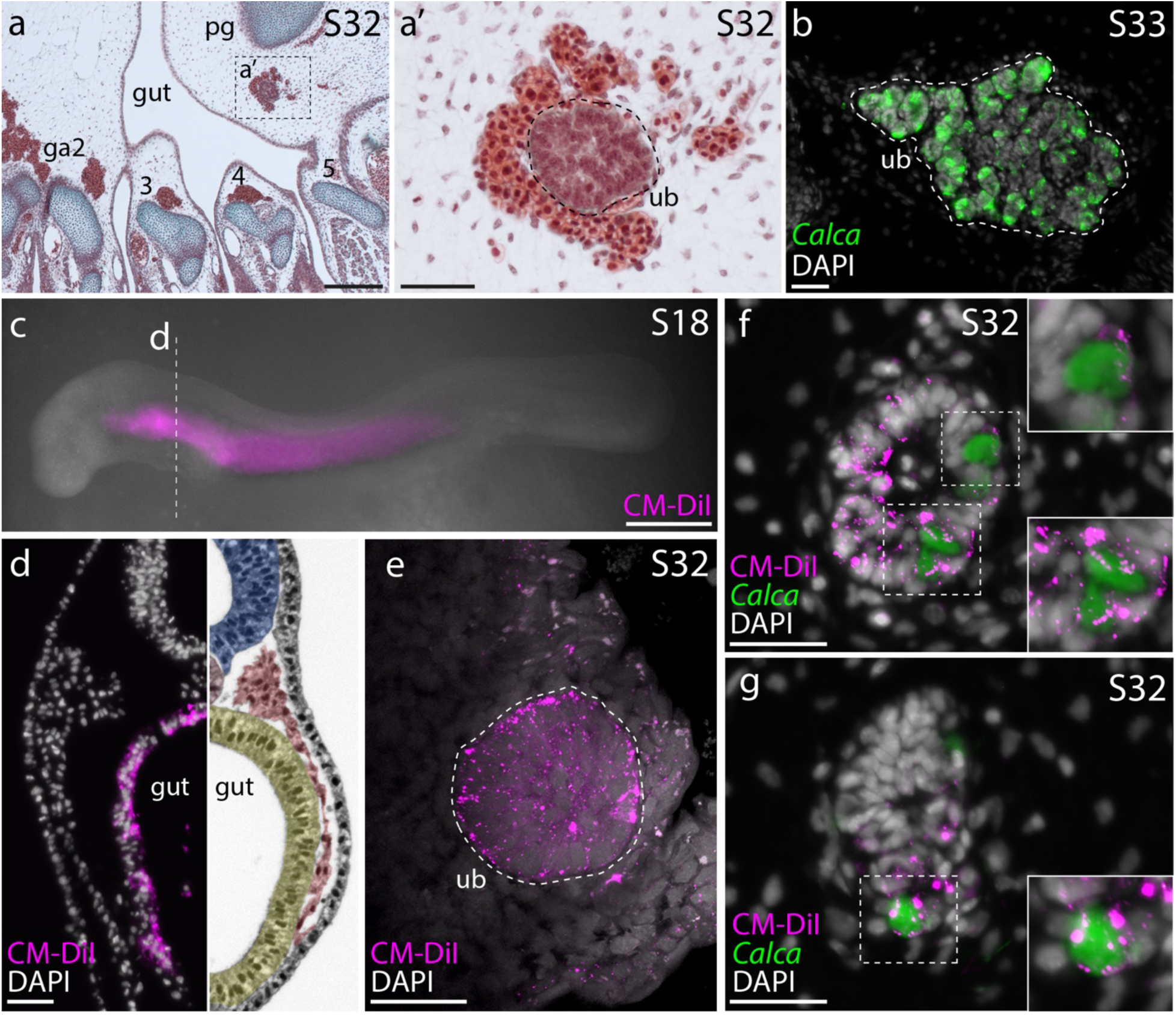
Skate C-cells derive from pharyngeal endoderm. **a,a’)** In a S32 skate embryo, the developing UB is located within connective tissue between the fifth gill arch and the pectoral girdle. **b)** The UB of a S33 skate embryo, with *Calca* expression in C-cells. **c)** Microinjection of CM-DiI into the pharyngeal cavity of a S18 skate embryo **d)** specifically labels the pharyngeal endoderm. **e)** Maximum intensity projection of the UB of a S32 skate embryo that received CM-DiI labeling of pharyngeal endoderm at S18. There is abundant CM-DiI labeling throughout the UB. **f, g)** Co-localisation of CM-DiI and *Calca* expression within the UB of S32 skate embryos indicate endodermal origin of C-cells in the skate. *ga2-5*, gill arches 2-5; *pg*, pectoral girdle; *ub*, ultimobranchial body. Scale bars: **a** = 200 μm; **a’** = 50 μm; **b** = 25 μm; **c** = 500 μm; **d,e,f,g** = 25 μm.

### Identification of putative endoderm-derived C-cells in non-vertebrate chordates

Tunicates and cephalochordates are invertebrate chordates that lack a *bona fide* neural crest, but that share many other bodyplan features with vertebrates, including an endodermally-derived pharynx^48^. Previous studies reported granulated and argyrophilic cell types in the endostyle of the tunicate *Styela clava*^49^ and in the stomach of the tunicate *Ciona intestinalis*^50^ that showed reactivity when stained with an anti-human calcitonin antibody. *C. intestinalis* has a single calcitonin-like gene (*Ci-CT*)^51–53^. We tested for the expression of *Ci-CT* in sections of the entire adult *C. intestinalis* using mRNA *in situ* hybridization by chain reaction (HCR). The adult life stage of *C. intestinalis* (Fig. 5a) possesses an endodermally-derived branchial sac^54^ that is perforated by numerous pharyngeal slits. This branchial sac is bordered on one side by an endostyle and on the other side by the gut (Fig. 5b) and bears inward-directed epithelial elaborations called papillae^55^. Consistent with previous studies^50^, we observed Ci-CT-expression in clusters of cells around the stomach of *C. intestinalis* (Fig. S3), though we additionally observed widespread expression in the lining of the pharynx (Fig. 5c). This pharyngeal expression localized to histologically distinct cells within the papillae of the branchial sac (Fig. 5d-e). These *Ci-CT*-expressing cells also report positively for the expression of *prohormone convertase 2*, an endopeptidase that converts prohormones and neuropeptide precursors to their active forms^55^, pointing to a likely neuroendocrine function for these cells. Contrary to previous studies^49,53^, we found no expression of *Ci-CT* in the endostyle of adult *C. intestinalis*.

**Figure 5:**
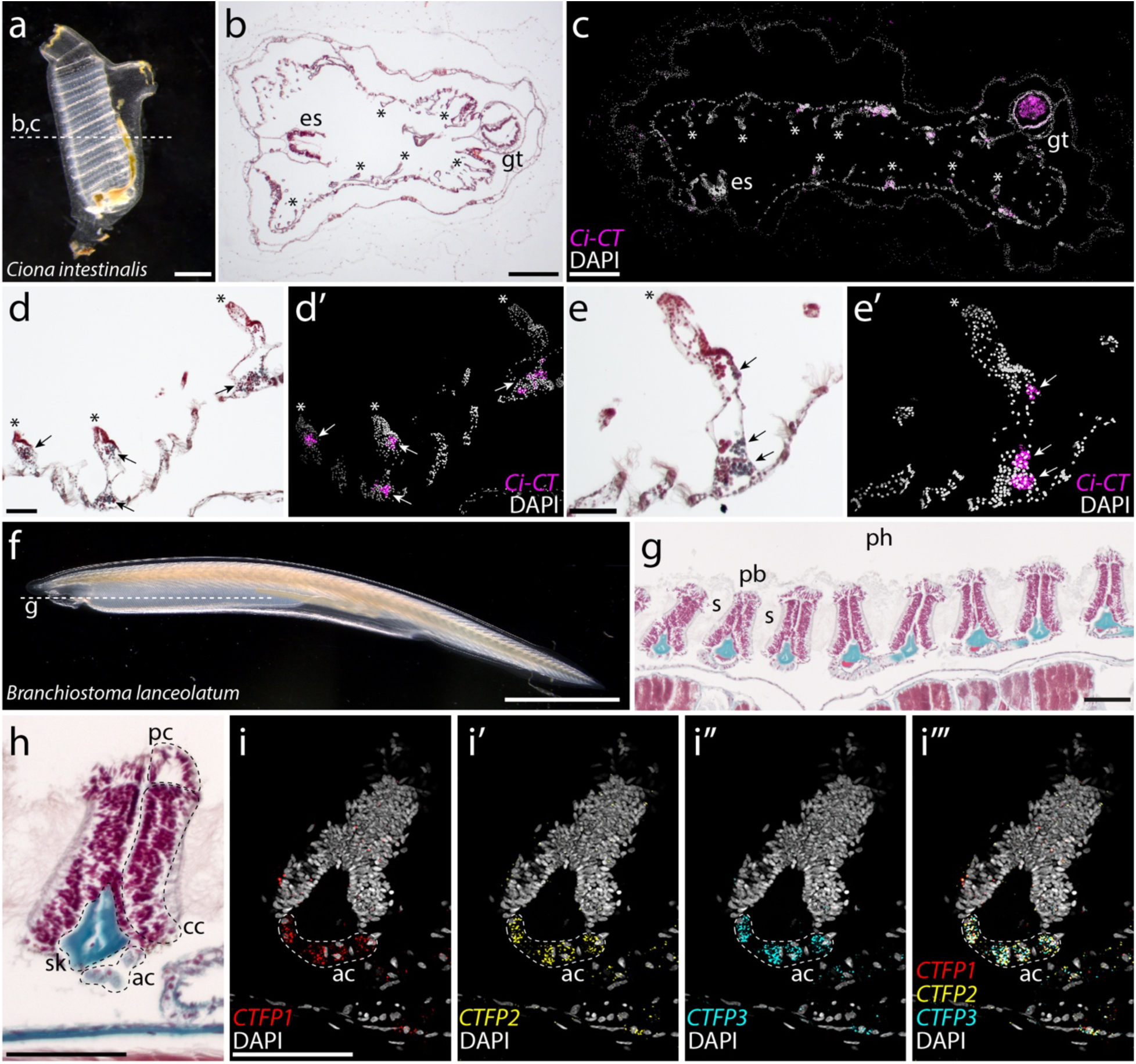
Putative C-cell homologues within the pharynx of invertebrate chordates. **a)** Adult *Ciona intestinalis*. Sections in the plane indicated by the dashed line reveal **b)** a branchial sac that is perforated by slits, with an endostyle on one side and the gut on the other. **c)** HCR for *Ci-CT* reveals expression in the endodermally-derived wall of the branchial sac (note autofluorescence within the gut). * in **b)** and **c)** indicate the papillae of the branchial sac. **d-e)** Expression of Ci-CT localizes to distinct clusters of cells within the papilla of the branchial sac. No expression was observed within the endostyle. **f)** An adult amphioxus, *Branchiostoma lanceolatum*. Sections in the plane indicated by the dashed line reveal **g)** a pharyngeal cavity containing a series of pharyngeal bars separated by pharyngeal slits. **h)** Each pharyngeal bar consists of atrial cells, ciliated cells, and pharyngeal cells, and is supported by an acellular skeletal rod. **i)** HCR for amphioxus *CTFP1-3* reveals co-expression of all three paralogues within the atrial cells of each pharyngeal bar. *ac*, atrial cells; *cc*, ciliated cells; *es*, endostyle; *gt*, gut; *pc*, pharyngeal cells; *sk*, skeletal rod. Scale bars: **a** = 1 mm; **b,c** = 250 μm; **d,e** = 50 μm; **f** = 2mm; **g,h,i** = 75 μm.

The pharynx of adult amphioxus (Fig. 5f) consists of a series of ∼50 pharyngeal slits that are separated by bars and is bordered ventrally by an endostyle (Fig. 5g). Pharyngeal bars have a thick surface epithelium divided into three portions: atrial cells that line the lateral surface of the bars, ciliated cells that are located on the medial side of each bar, and pharyngeal cells that line the bar between the atrial and ciliated cells (Fig. 5h). Pharyngeal bars are also supported internally by a skeletal rod, consisting of an expanded acellular collagenous matrix^56^. Cephalochordates have three calcitonin genes, called *CTFP*s (calcitonin family proteins), that are expressed in different combinations in neurons of the amphioxus larva^57–59^. We identified the complete sequence and genomic location of the three CTFP genes in *Branchiostoma lanceolatum*. Phylogenetic analysis and the close arrangement of *CTFP* genes on the same chromosome together indicate that *CTFP*s duplicated independently in the cephalochordate lineage (Fig. S4). We generated probes for all three *CTFP* paralogs and tested for their expression in longitudinal sections of the adult *Branchiostoma lanceolatum* pharynx using *in situ* HCR. We found co-expression of all three *CTFP* paralogs in the atrial cells of each pharyngeal bar (Fig. 5i; Fig. S5), and we detected no expression of any *CTFP* paralogue in the endostyle.

These findings point to a calcitonin-expressing cell type within the endodermally-derived pharyngeal lining of tunicates and cephalochordates, and to an ancient, pre-vertebrate endodermal origin of calcitonin-secreting neuroendocrine cells. There is deep conservation across deuterostomes of a core pharyngeal endodermal transcriptional program^60^, and genes encoding transcription factors like Pax1/9^61^, Six1^62^ and Eya1^63^ that specify glandular tissues (e.g., the thymus, parathyroid and UBs) within the caudal pharyngeal endodermal pouches of vertebrate embryos are expressed iteratively in all developing pharyngeal pores of amphioxus^64,65^ and hemichordates^66^. The iterative deployment of this conserved transcriptional program throughout pharyngeal development could account for the broader distribution of some neuroendocrine cell types, such as C-cells, within pharyngeal endodermal derivatives of invertebrate chordates, with these cells subsequently becoming localized to specific glandular structures with tissue elaboration and enhanced regionalization of the vertebrate pharynx (Fig. 6).

**Figure 6:**
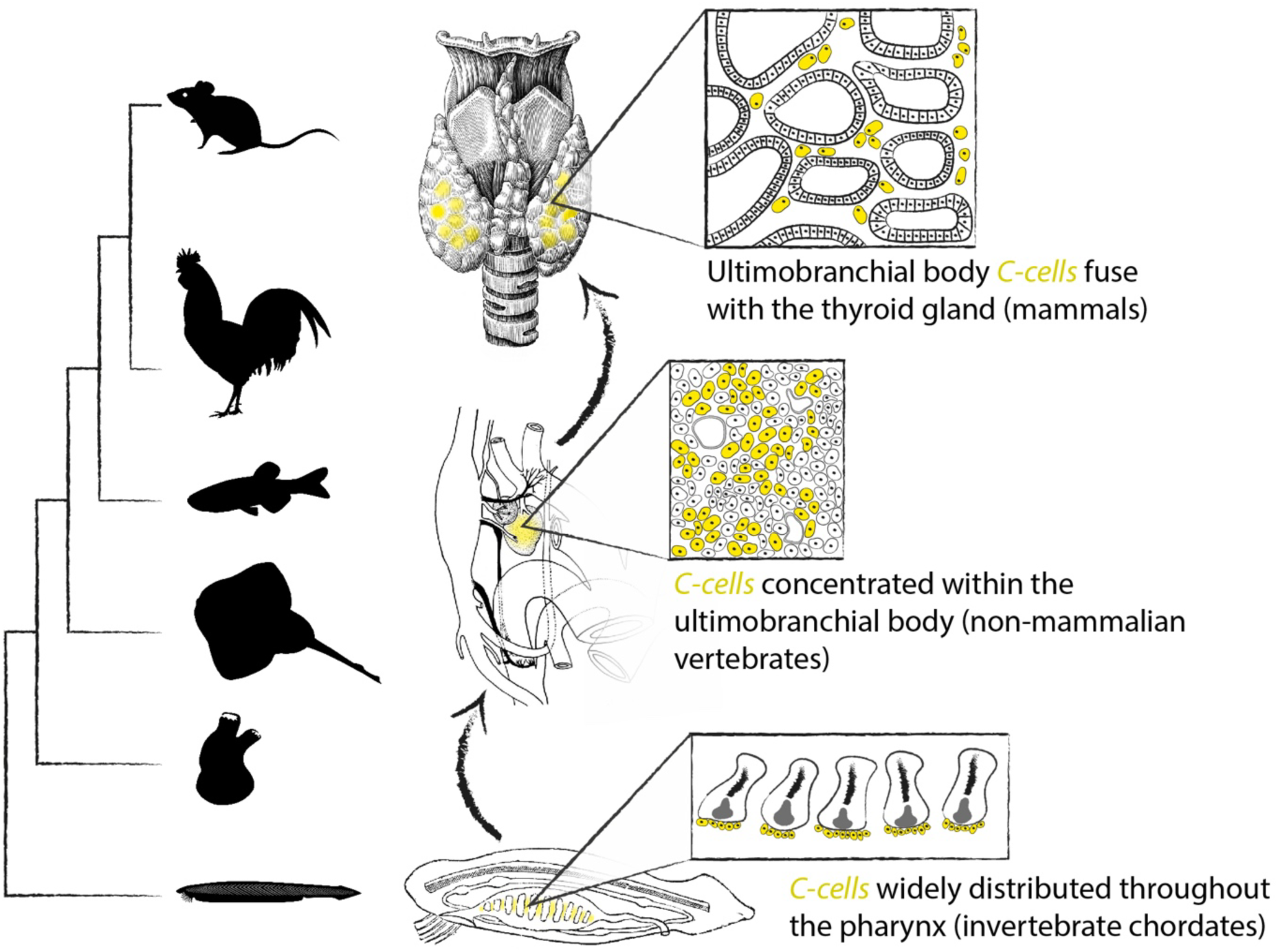
Evolution and homology of chordate C-cells. C-cells were broadly distributed throughout the endoderm-derived lining of the pharynx in the last common ancestor of chordates. These C-cells then became localized into a discrete ultimobranchial body in non-mammalian vertebrates, with that ultimobranchial body fusing with the thyroid gland of mammals.

## Conclusion

Here, we report evidence of a broadly conserved endodermal origin of C-cells in tetrapods, bony fishes, and cartilaginous fishes. This stands in contrast to the previously reported neural crest origin of avian C-cells, and points instead to an ancestral endodermal origin of this cell type. Additionally, we report a discrete calcitonin-positive cell type (a putative C-cell homologue) that resides within the endodermally-derived pharyngeal lining of the invertebrate chordates *Ciona intestinalis* and *Branchiostoma lanceolatum*. These observations support a pre-vertebrate origin of an endodermally-derived calcitonin-expressing neuroendocrine cell, broaden the ancestral chordate endodermal cell type repertoire to include a new neuroendocrine lineage, and establish homology of C-cells throughout chordate phylogeny.

## Materials and Methods

### Zebrafish lines

The following zebrafish (*Danio rerio*) lines were used: Tg(Mmu.Sox10-Mmu.Fos:Cre)^zf384Tg^ ^41^, Tg(actab2:loxP-BFP-STOP-loxP-dsRed)^sd2Tg^ ^67^, Tg(-5.0sox17:creERT2; myl7:DsRed)^sid1Tg^ ^68^, Tg(-3.5ubb:loxP-eGFPloxP-mCherry)^cz1701Tg^ ^69^ and *tfap2a^mob^;foxd3^mos^* ^40^. To induce Cre recombination in the endoderm, Tg(-5.0sox17:creERT2;myl7:DsRed)^sid1Tg^; Tg(-3.5ubb:loxP-GFPloxP-mCherry)^cz1701Tg^ embryos were incubated in embryo media containing 5 μM (Z)−4-Hydroxytamoxifen (Sigma-Aldrich H7904) starting at 4-4.5 hours post-fertilization (hpf) and then washed several times in fresh embryo media at 6 hpf. For lineage tracing experiments, only preselected animals with high conversion were used. Experiments using all zebrafish lines were conducted according to protocols approved by the Institutional Animal Care and Use Committees in facilities accredited by the Association for Assessment and Accreditation of Laboratory Animal Care International (AAALAC). All zebrafish were fixed overnight at 4°C in 4% paraformaldehyde in phosphate-buffered saline (PBS), rinsed in PBS, and dehydrated into methanol prior to analysis.

### Skate embryos

Skate (*L. erinacea*) embryos were obtained from the Marine Resources Center at the Marine Biological Laboratory (MBL) in Woods Hole, MA, U.S.A., were reared and stages as described in Gillis *et al.* (2022)^44^. All skate experiments were conducted according to protocols approved by the Institutional Animal Care and Use Committee of the MBL. Endodermal lineage tracing was performed by microinjection of CellTracker CM-DiI (1,1’-dioctadecyl-3,3,3’3’-tetramethylindocarbocyanine perchlorate) into the pharyngeal cavity of S18 skate embryos with a pulled glass needle. CM-DiI was prepared as previously described^70^. Skate embryos were euthanized with an overdose of MS-222 (1 g/L in seawater), and all embryos were fixed overnight at 4°C in 4% paraformaldehyde in phosphate-buffered saline PBS, and then rinsed in PBS and dehydrated into methanol prior to analysis.

### Chicken embryos

Experiments using chicken (*Gallus gallus domesticus*) embryos were conducted in accordance with the UK Animals (Scientific Procedures) Act 1986. Fertilised wild-type chicken eggs were obtained from commercial sources. Fertilised GFP-transgenic chicken eggs^39^ were obtained from the Roslin Institute Transgenic Chicken Facility (Edinburgh, UK), which is funded by Wellcome and the BBSRC. Wild-type and GFP-transgenic eggs were incubated in a humidified atmosphere at 38 °C for approximately 1.5 days to reach 6–11 somites and embryos visualised as previously described^71^, using filtered PBS instead of Ringer’s solution. To label premigratory vagal neural crest cells, neural fold between the level of the otic vesicle and the caudal end of (1) somite 1 (s1) (unilaterally) or (2) s6 (unilaterally) was grafted isotopically from GFP-transgenic donors to wild-type hosts using a pulled glass needle. Chick embryos were fixed overnight at 4°C in 4% paraformaldehyde in phosphate-buffered saline PBS, and then rinsed in PBS and dehydrated into methanol prior to analysis.

### Branchiostoma and Ciona collection

Adults of the European amphioxus (*Branchiostoma lanceolatum*) were collected in Banyuls-sur-Mer, France and maintained in a custom-made facility at the Department of Zoology, University of Cambridge^72^. Animals were anesthetized with a 30 min immersion in 0.015% tricaine methanesulphonate (MS-222), divided into rostral and caudal halves using a razor blade, and fixed in 4% paraformaldehyde for 24 hours at 4°C, as described^73^. Adults of the sea squirt *Ciona intestinalis* were kept in tanks within the amphioxus facility at the Department of Zoology, University of Cambridge. Small adult ascidians were detached from the tanks, relaxed with menthol crystals dissolved in seawater and fixed in 4% paraformaldehyde for 24 hours at 4°C. The anterior half of fixed amphioxus and whole *Ciona* young adults were then processed for sectioning and *in situ* hybridization.

### Sequence analysis

The complete prepropeptide sequences of multiple vertebrate calcitonin and calcitonin gene-related peptides, as well as of *Ciona*, amphioxus, and starfish calcitonin-type prepropeptides (Table S1) were aligned using MAFFT^74^ and trimmed using trimAl^75^. Neighbor joining molecular phylogenetics was carried out with seaview^76^, using the related neuropeptide adrenomedullin as outgroup, and the resulting tree visualized with FigTree (http://tree.bio.ed.ac.uk/software/figtree/).

### Sectioning and staining

Prior to embedding, adult zebrafish were decalcified in Morse solution (10% w/v sodium citrate dihydrate and 25% v/v formic acid in DEPC water) for 24 hours at room temp with gentle agitation. All tissue samples were cleared with Histosol (National Diagnostics) for 3 x 20 min at room temperature, transitioned into 1:1 Histosol:Paraffin for 2 x 30 min at 60°C, then infiltrated with molten paraffin overnight at 60°C. After an additional 4 x 1 hour paraffin changes, samples were embedded in peel-a-way molds (Sigma), left to set for 24 hours and then sectioned at 7 μm on a Leica RM2125 rotary microtome. Sections were mounted on SuperFrost Plus charged glass slides. Histochemical staining with modified Masson’s Trichrome was performed as described by Witten and Hall^77^.

### mRNA *in situ* hybridization on paraffin sections

Chromogenic mRNA *in situ* hybridization (ISH) was performed on paraffin sections as previously described^78^ with modifications^70^. Chromogenic ISH probes against zebrafish *calca* (GenBank DQ406589.1) and skate *Calca* (GenBank XM_055649884.1) were generated by *in vitro* transcription using standard methods. Third-generation mRNA ISH by chain reaction (HCR) was performed as per Choi et al.^79^ following the protocol for formaldehyde-fixed, paraffin-embedded sections, with modifications^80^. For chick and skate, probes, buffers, and hairpins were purchased from Molecular Instruments (Los Angeles, California, USA). HCR probe set lot numbers are as follows: Chicken *Calca* (###), Chicken *TH* (###) and Skate *Calca* (###). Ciona Ci-CT (###), and Branchiostoma CTFP1 (###), CTFP2 (###) and CTFP3 (###).

### Immunofluorescence on paraffin sections

All slides for immunofluorescence were dewaxed for 2 x 5min in Histosol, rehydrated through a descending ethanol series, and washed 3 x 5 min in PBS + 0.1% Triton X-100 (PBST). Antigen retrieval was performed by pre-warming slides in distilled water at 60°C for 5 min, then incubating in 10mM tri-sodium citrate (pH6.0) at 95°C for 25 min. Slides were then cooled at -20°C for 30 min and rinsed 3 x 5 min in PBST before blocking in 10% heat-inactivated sheep serum at room temperature for 60 min. Primary antibodies were applied underneath a parafilm coverslip, and slides were incubated in a humidified chamber overnight at 4°C. Following primary antibody incubation, slides were rinsed 3 x 10 min with PBST, and then secondary antibodies were applied underneath a parafilm coverslip and slides were incubated at room temperature for 4 hours in the dark. Slides were then rinsed 3 x 10 min in PBST, then 3 X 30 min in PBST before coverslipping with Fluoromount G containing DAPI. Primary antibodies used were: mouse anti-mCherry (Abcam, ab125096; 1:250), mouse anti-GFP (Merck, SAB5300167; 1:250), rabbit anti-tyrosine hydroxylase (Merck, AB152; 1:250), rabbit-anti calcitonin (BMA Biomedicals, T-4026; 1:250), Goat anti-Mouse IgG (H+L) Cross-Adsorbed Secondary Antibody, Alexa Fluor 488 (ThermoFisher, A11001; 1:500), Goat anti-Rabbit IgG (H+L) Cross-Adsorbed Secondary Antibody, Alexa Fluor 488 (ThermoFisher, A11008; 1:500), Goat anti-Mouse IgG (H+L) Cross-Adsorbed Secondary Antibody, Alexa Fluor 633 (ThermoFisher, A21050; 1:500) and Goat anti-Rabbit IgG (H+L) Highly Cross-Adsorbed Secondary Antibody, Alexa Fluor 633 (ThermoFisher, A21071; 1:500).

### Imaging and image processing

Images were taken on a Zeiss Axioscope A1 compound microscope with a Zeiss Colibri 7 fluorescence LED light source using a Zeiss Axiocam 305 color or 503 mono camera and ZenPro software. All figures were assembled using Adobe creative cloud. Images of chromogenic ISH were inverted and overlaid with corresponding DAPI images.

## Acknowledgments

We would like to thank Prof. Clare Baker and Prof. Roger Keynes for helpful advice throughout this study. This work was supported by Wellcome Ph.D. Studentships to J.M.R. (214953/Z/18/Z) and D.H. (086804/Z/08/Z); by a Whitten Studenship to G.G.; by NIMH (R01MH113362) to E.W.K.; by NIGMS (T32GM008554) and NIDCR (F31DE030007) to D.J.R; by CRUK (C9545/A29580) to E.B.G; by NIDCR (R35DE027550) to J.G.C.; by NIDCR K99 DE029858 and GACR 23-06977M to P.F.; and by a Royal Society University Research Fellowship (UF130182 and URF\R\191007), Royal Society Research Grant (RG140377) and Marine Biological Laboratory Institutional Funding to J.A.G.

## Author contributions

J.A.G. conceived and designed the study; J.A.G. collected and analysed all zebrafish gene expression and lineage tracing data, collected all chick embryo gene expression data, analysed chick embryo fate mapping experiments, and performed all skate CM-DiI lineage tracing experiments; J.M.R. performed and analysed all chick embryo endodermal lineage tracing experiments; K.K. analysed skate lineage tracing experiments; G.G. prepared, imaged and analysed all *Ciona* and *Branchiostoma* gene expression data; D.H. performed all chick neural tube transplantation experiments; D.J.R. and EWK provided fixed *tfap2a^mob^;foxd3^mos^* zebrafish; J.G.C. and P.F. provided fixed sox17:CreERT2; ubi:loxP-eGFP-STOP-loxP-mCherry and Sox10:Cre; actab2:loxP-BFP-STOP-loxP-dsRed (Sox10>dsRed) fish; J.A.G. supervised the project with support from E.B.-G.; J.A.G. wrote the manuscript and prepared all figures, with input from all co-authors.

## Declaration of interests

EBG has been employed by Genentech since September 2022. The rest of the authors declare no competing interests.

## References

1. Copp, D.H., and Cheney, B. (1962) Calcitonin—a hormone from the parathyroid which lowers the calcium-level of the blood. Nature 193, 381–382.

2. Copp, D.H., Cameron, E.C., Cheney, B.A., Davidson, A.G., and Henze, K.G. (1962) Evidence for calcitonin—a new hormone from the parathyroid gland that lowers blood calcium. Endocrinology 70, 638–649.

3. Pearse, A.G. (1966) The cytochemistry of the thyroid C cells and their relationship to calcitonin. Proc R Soc London B Bio Sci 164, 478–487.

4. Pearse, A.G. (1966) 5-Hydroxytryptophan uptake by dog thyroid ‘C’ cells, and its possible significance in polypeptide hormone production. Nature 211, 598–600.

5. Chambers, T.J., and Moore A. (1983) The sensitivity of isolated osteoclasts to morphological transformation by calcitonin. J Clin Endocrinol Metab 57, 819–824.

6. Nicholson, G.C., Moseley, J.M., Sexton, P.M., Mendelsohn F.A., and Martin, T.J. (1986) Abundant calcitonin receptors in isolated rat osteoclasts. Biochemical and autoradiographic characterization. J Clin Invest 78, 355– 359.

7. Talmage, R.V., Grubb, S.A., Norimatsu, H., and Vanderwiel, C.J. (1980) Evidence for an important physiological role for calcitonin. Proc Natl Acad Sci USA 77, 609–613.

8. Muñoz-Torres, M., Alonso, G., and Raya, M.P. (2004) Calcitonin therapy in osteoporosis. Treat Endocrinol 3, 117– 132.

9. Langston, A.L., and Ralston, S.H. (2004) Management of Paget’s disease of bone. Rheumatology 43, 955–959.

10. Hurley, D.L., Tiegs, R.D., Wahner, H.W., Heath 3rd, H. (1987) Axial and appendicular bone mineral density in patients with long-term deficiency or excess of calcitonin. N Engl J Med 317, 537–541.

11. Wuster, C., Raue, F., Meyer, C., Bergmann, M., and Ziegler, R. (1992) Long-term excess of endogenous calcitonin in patients with medullary thyroid carcinoma does not affect bone mineral density. J Endocrinol 134, 141–147.

12. Zaidi, M., Fuller, K., Bevis, P.J., GainesDas, R.E., Chambers, T.J., and MacIntyre, I. (1987) Calcitonin gene-related peptide inhibits osteoclastic bone resorption: a comparative study. Calcif Tissue Int 40, 149–154.

13. Hoff, A.O., et al. (2002) Increased bone mass is an unexpected phenotype associated with deletion of the calcitonin gene. J Clin Invest 110, 1849–1857.

14. Janine, P. et al. (2006) Calcitonin plays a critical role in regulating skeletal mineral metabolism during lactation. Endocrinology 147, 4010–4021.

15. Srivastav, A.K., Srivastav, S.K., Sasayama, Y., and Suzuki, N. (1998) Salmon calcitonin induced hypocalcemia and hyperphosphatemia in an elasmobranch, *Dasyatis akajei*. Gen Comp Endocrinol 109, 8–12.

16. Glowacki, J., O’Sullivan, J., Miller, M., Wilkie, D.W., and Deftos, L.J. (1985) Calcitonin produces hypercalcemia in leopard sharks. Endocrinology 116, 827–829.

17. Suzuki, N., Takagi, T. Sasayama, Y., and Kambegawa, A. (2009). Effects of ultimobranchialectomy on the mineral balances of the plasma and bile in the stingray (Elasmobranchii). Zool Sci 12, 239–242.

18. Cardoso, J.C.R., Félix, R.C., Ferreira, V., Peng, M., Zhang, X., and Power, D.M. (2020) The calcitonin-like system is an ancient regulatory system of biomineralization. Sci Rep 10, 7581.

19. Dean, M.N., Ekstrom, L., Monsonego-Ornan, E., Ballantyne, J., Witten, P.E., Riley, C., Habraken, W., and Omelon, S. (2015) Mineral homeostasis and regulation of mineralization processes in the skeletons of sharks, rays and relatives (Elasmobranchii). Semin Cell Dev Biol 46, 51–67.

20. Hamilton, W.J., Boyd, J.D., and Mossman, H.W. (1945) Human Embryology (W. Heffer & Sons Limited, Cambridge).

21. Nilsson, M., and Fagman, H. (2017) Development of the thyroid gland. Development 144, 2123–2140.

22. Tauber, S.D. (1967) The ultimobranchial origin of thyrocalcitonin. Proc Natl Acad Sci USA 58, 1684–1687.

23. Copp, D.H., Cockcroft, D.W., and Kueh, Y. (1967) Calcitonin from ultimobranchial glands of dogfish and chickens. Science 158, 924–925.

24. Kameda, Y. (2017) Morphological and molecular evolution of the ultimobranchial gland of nonmammalian vertebrates, with special reference to the chicken C cells. Dev Dyn 246, 719–739.

25. 25. Le Douarin, N., and Le Lièvre, C. (1970) Démonstration de l’origine neurale des cellules à calcitonine de corps ultimobranchial chez l’embryon de poulet. C R Acad Sc Paris 270, 2875–2860.

26. Pearse, A.G., and Polak, J.M. (1971) Cytochemical evidence for the neural crest origin of mammalian ultimobranchial C cells. Histochemie 27, 96–102.

27. 27. Polak, J.M., Pearse, A.G., Le Lièvre, C., Fontaine, J., and Le Douarin, N.M. (1974) Immunocytochemical confirmation of the neural crest origin of avian calcitonin-producing cells. Histochemistry 40, 209–214.

28. 28. Master, S.R., and Burns, B. (2023) Medullary Thyroid Cancer. 2023 Feb 15. In: StatPearls. Treasure Island (FL): StatPearls Publishing. PMID: 29083765.

29. Segura, S., Ramos-Rivera, G., and Suhrland, M. (2018) Educational Case: Endocrine Neoplasm: Medullary Thyroid Carcinoma. Acad Pathol 5, 2374289518775722.

30. Gagel, R.F., and Cote, G.J. (1998) Pathogenesis of Medullary Thyroid Carcinoma. In: Fagin, J.A. (eds) Thyroid Cancer. Endocrine Updates, vol 2 (Springer, Boston).

31. Pearse, A.G. (1969) The cytochemistry and ultrastructure of polypeptide hormone-producing cells of the APUD series and the embryologic, physiologic and pathologic implications of the concept. J Histochem Cytochem 17, 303–313.

32. Pearse, A.G., and Polak, J.M. (1974) Endocrine tumours of neural crest origin: neurolophomas, apudomas and the APUD concept. Med Biol 52, 3–18.

33. Pearse, A.G., and Polak, J.M. (1971) Neural crest origin of the endocrine polypeptide (APUD) cells of the gastrointestinal tract and pancreas. Gut 12, 783–788.

34. Kameda, Y., Nishimaki, T., Chisaka, O., Iseki, S., and Sucov, H.M. (2007) Expression of the epithelial marker E-cadherin by thyroid C cells and their precursors during murine development. J Histochem Cytochem 55, 1075– 1088.

35. Johansson, E., et al. (2015) Revising the embryonic origin of thyroid C cells in mice and humans. Development 142, 3519–3528.

36. Andrew, A., Kramer, B., and Rawdon, B.B. (1983) Gut and pancreatic amine precursor uptake and decarboxylation cells are not neural crest derivatives. Gastroenterology 84, 429–431.

37. Kuo, C.S., and Krasnow, M.A. (2015) Formation of a neurosensory organ by epithelial cell slithering. Cell 163, 394– 405.

38. Andrew, A. (1974) Further evidence that enterochromaffin cells are not derived from the neural crest. J Embryol Exp Morphol. 31, 589–598.

39. McGrew, M.J. et al. (2008) Localised axial progenitor cell populations in the avian tail bud are not committed to a posterior Hox identity. Development 135, 2289–2299.

40. Wang, W.-D., Melville, D.B. Montero-Balaguer, M., Hatzopoulos, A.K., and Knapik, E.W. (2011) Tfap2a and Foxd3 regulate early steps in the development of the neural crest progenitor population. Dev Biol 360, 173–185.

41. Kague, E., Gallagher, M., Burke, S., Parsons, M., Franz-Odendaal, T., and Fisher, S. (2012) Skeletogenic fate of zebrafish cranial and trunk neural crest. PLoS One 7, e47394.

42. Fabian, P., Tseng, K.C., Thiruppathy, M., Arata, C., Chen, H.J., Smeeton, J., Nelson, N., and Crump, J.G. (2022) Lifelong single-cell profiling of cranial neural crest diversification in zebrafish. Nat Commun 13, 13.

43. Fabian, P., Tseng, K.C., Smeeton, J., Lancman, J.J., Dong, P.D.S., Cerny, R., and Crump, J.G. (2020) Lineage analysis reveals an endodermal contribution to the vertebrate pituitary. Science 370, 463–467.

44. Gillis, J.A., Bennett, S., Criswell, K.E., Rees, J., Sleight, V.A., Hirschberger, C., Calzarette, D., Kerr, S., and Dasen, J. (2022) Big insight from the little skate: *Leucoraja erinacea* as a developmental model system. Curr Top Dev Biol 147, 595–630.

45. Gillis, J.A., and Tidswell, O.R. (2017) The origin of vertebrate gills. Curr Biol 27, 729–732.

46. Rees, J.M., Sleight, V.A., Clark, S.J., Nakamura, T., and Gillis, J.A. (2023) Ectodermal Wnt signaling, cell fate determination, and polarity of the skate gill arch skeleton. Elife 12, e79964.

47. Sleight, V.A., and Gillis, J.A. (2020) Embryonic origin and serial homology of gill arches and paired fins in the skate, *Leucoraja erinacea*. eLife 9, e60635.

48. Lowe, C.J., Clarke, D.N., Medeiros, D.M., Rokhsar, D.S., and Gerhart, J. (2015) The deuterostome context of chordate origins. Nature 520, 456–465.

49. Thorndyke, M.C., and Probert, L. (1979) Calcitonin-like cells in the pharynx of the ascidian *Styela clava*. Cell Tissue Res 203, 301–309.

50. Fritsch, H.A., Van Noorden, S., and Pearse, A.G. Calcitonin-like immunochemical staining in the alimentary tract of *Ciona intestinalis* L. Cell Tissue Res 205, 439–444.

51. Hamada, M., Shimozono, N., Ohta, N., Satou, Y., Horie, T., Kawada, T., Satake, H., Sasakura, Y., and Satoh, N. (2011) Expression of neuropeptide- and hormone-encoding genes in the *Ciona intestinalis* larval brain. Dev Biol 352, 202–214.

52. Satake, H., and Sasakura, Y. (2023) The neuroendocrine system of *Ciona intestinalis* Type A, a deuterostome invertebrate and the closest relative of vertebrates. Mol Cell Endocrinol 582, 112122.

53. Sekiguchi T et al. (2009) Calcitonin in a protochordate, *Ciona intestinalis* -- the prototype of the vertebrate calcitonin/calcitonin gene-related peptide superfamily. FEBS J 276, 4437–4447.

54. Hirano, T., and Nishida, H. (2000) Developmental fates of larval tissues after metamorphosis in the ascidian, Halocynthia roretzi. II. Origin of endodermal tissues of the juvenile. Dev Genes Evol 210, 55–63.

55. Osugi, T., Sasakura, Y., and Satake, H. (2020) The ventral peptidergic system of the adult ascidian *Ciona robusta* (*Ciona intestinalis* Type A) insights from a transgenic animal model. Sci Rep 10, 1892.

56. Baskin, D.G., and Detmers, P.A. (1976) Electron microscopic study on the gill bars of amphioxus (*Branchiostoma californiense*) with special reference to neurociliary control. Cell Tissue Res 166, 167–178.

57. Sekiguchi, T. (2018) The calcitonin/calcitonin gene-related peptide family in invertebrate deuterostomes. Front Endocrinol 9, 695.

58. Sekiguchi T et al. (2016) Evidence for conservation of the calcitonin superfamily and activity-regulating mechanisms in the basal chordate *Branchiostoma floridae*: Insights into the molecular and functional evolution in chordates. J Biol Chem 291, 2345–2356.

59. 59. Gattoni, G., Keitley, D., Sawle, A., and Benito-Gutiérrez, E. (2023) An ancient gene regulatory network sets the position of the forebrain in chordates bioRxiv 2023.03.13.532359. doi: 10.1101/2023.03.13.532359

60. Simakov, O. et al. (2015) Hemichordate genomes and deuterostome origins. Nature 527, 459–465.

61. Peters, H., Neubüser, A., Kratochwil, K., and Balling, R. (1998) Pax9-deficient mice lack pharyngeal pouch derivatives and teeth and exhibit craniofacial and limb abnormalities. Genes Dev 12:2735–2747.

62. Zou, D., Silvius, D., Davenport, J., Grifone, R., Maire, P., and Xu, P.X. (2006) Patterning of the third pharyngeal pouch into thymus/parathyroid by Six and Eya1. Dev Biol. 293, 499–512.

63. Xu, P.X., Zheng, W., Laclef, C., Maire, P., Maas, R.L., Peters, H., and Xu, X. (2002) Eya1 is required for the morphogenesis of mammalian thymus, parathyroid and thyroid. Development 129, 3033 – 3044.

64. Liu, X., Li, G., Liu, X., and Wang, Y.Q. (2015) The role of the Pax1/9 gene in the early development of amphioxus pharyngeal gill slits. J Exp Zool B Mol Dev Evol 324, 30–40.

65. Kozmik, Z. et al. (2007) Pax-Six-Eya-Dach network during amphioxus development: conservation in vitro but context specificity in vivo. Dev Biol 306, 143–159.

66. Gillis, J.A., Fritzenwanker, J.H., and Lowe, C.J. A stem-deuterostome origin of the vertebrate pharyngeal transcriptional network. Proc Biol Sci 279, 237–246.

67. Kobayashi, I., Kobayashi-Sun, J., Kim, A.D., Pouget, C., Fujita, N., Suda, T., and Traver, D. (2014) Jam1a–Jam2a interactions regulate haematopoietic stem cell fate through Notch signalling. Nature 512, 319–323.

68. Hockman, D. et al. (2017) Evolution of hypoxia-sensitive cells involved in amniote respiratory reflexes. eLife 6, e21231.

69. Mosimann, C., Kaufman, C.K., Li, P., Pugach, E.K., Tamplin, O.J., and Zon, L.I. (2011) Ubiquitous transgene expression and Cre-based recombination driven by the *ubiquitin* promoter in zebrafish. Development 138, 169– 177.

70. Gillis, J.A., Modrell, M.S., Northcutt, R.G., Catania, K.C., Luer, C.A., and Baker, C.V.H. (2012) Electrosensory ampullary organs are derived from lateral line placodes in cartilaginous fishes. Development 139, 3142–3146.

71. Dude, C.M., Kuan, C.-Y.K., Bradshaw, J.R., Greene, N.D.E., Relaix, F., Stark, M.R., and Baker C.V.H. (2009) Activation of Pax3 target genes is necessary but not sufficient for neurogenesis in the ophthalmic trigeminal placode. Dev Biol 326, 314–326.

72. Benito-Gutierrez, È., Weber, H., Bryant, D.V., and Arendt, D. (2013) Methods for generating year-round access to amphioxus in the laboratory. PLoS One 8, e71599.

73. Andrews, T.G.R., Gattoni, G., Busby, L., Schwimmer, M.A., and Benito-Gutierrez, È. (2020) Hybridization chain reaction for quantitative and multiplex imaging of gene expression in amphioxus embryos and adult tissues. Methods Mol Biol 2148, 179–194.

74. Katoh, K. and Standley, D.M. (2013) MAFFT Multiple Sequence Alignment Software Version 7: Improvements in Performance and Usability. Mol Biol Evol 30, 772–780.

75. Capella-Gutiérrez, S., Silla-Martínez, J.M., and Gabaldón, T. (2009) trimAl: a tool for automated alignment trimming in large-scale phylogenetic analyses. Bioinformatics 25, 1972–1973.

76. Gouy, M., Tannier, E., Comte, N., and Parsons, D.P. (2021) Seaview Version 5: A Multiplatform Software for Multiple Sequence Alignment, Molecular Phylogenetic Analyses, and Tree Reconciliation. Methods Mol Biol 2231, 241– 260.

77. Witten, P.E., and Hall, B.K. (2003) Seasonal changes in the lower jaw skeleton in male Atlantic salmon (*Salmo salar* L.): remodelling and regression of the kype after spawning. J Anat 203, 435–450.

78. O’Neill, P., McCole, R.B., Baker, C.V.H. (2007) A molecular analysis of neurogenic placode and cranial sensory ganglion development in the shark, *Scyliorhinus canicula*. Dev Biol 304, 156–181.

79. Choi, H.M.T., Schwarzkopf, M., Fornace, M.E., Acharya, A., Artavanis, G., Stegmaier, J., Cunha, A., and Pierce, N.A. (2018) Third-generation *in situ* hybridization chain reaction: multiplexed, quantitative, sensitive, versatile, robust. Development 145, dev165753.

80. Criswell, K.E., and Gillis, J.A. (2020) Resegmentation is an ancestral feature of the gnathostome vertebral skeleton. eLife 9, e51696.

