## Supplemental Figures for "A pre-vertebrate endodermal origin of calcitonin-producing neuroendocrine cells"

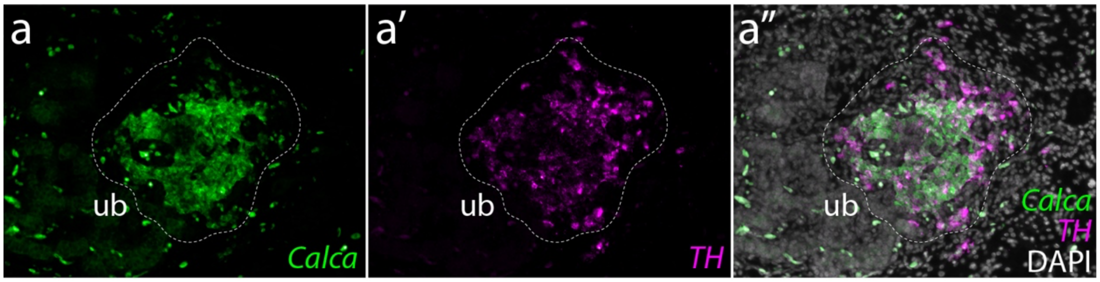


**Figure S1: Chick C-cells co-express *Calca* and *TH*. a)** *Calca* and **a')** *TH* are **a")** largely co-expressed in the C-cells of the chick UB. Expression detected by multiplexed fluorescent mRNA *in situ* hybridization by chain reaction. *ub*, ultimobranchial body.

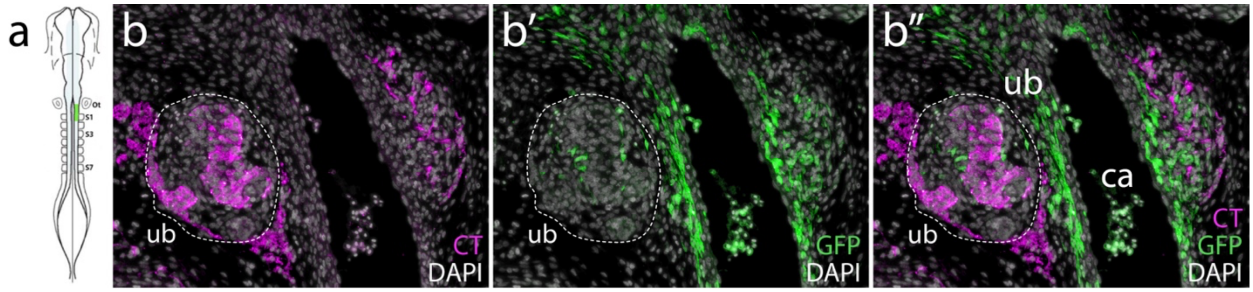


**Figure S2: Chick C-cells do not derive from the neural crest.** Neural crest lineage tracing was performed by isotopic unilateral grafting of GFP+ neural fold into a wild-type host embryo. **a)** In this example, a graft was performed with neural fold from the otic vesicle to S1. **b)** Calcitonin (CT) is expressed in the C-cells of the UBU. **b')** GFP+ neural crest derived cells are recovered within and around the UB, but **b")** we observed no co-localisation of GFP and CT. *ub*, ultimobranchial body.

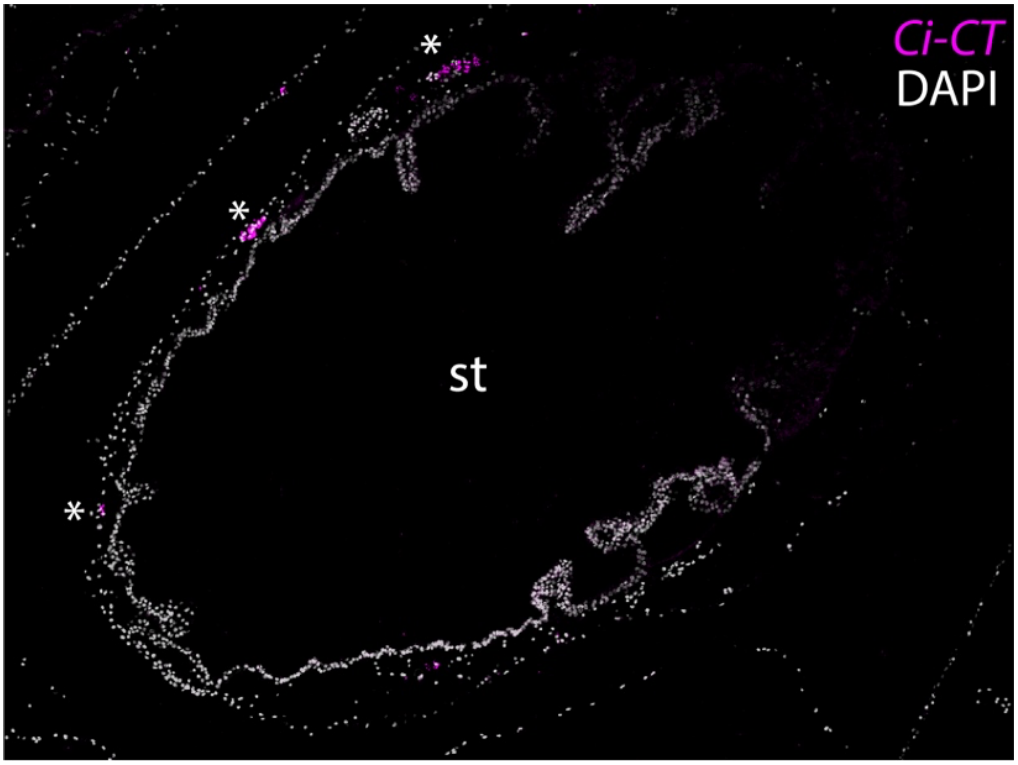


**Figure S3: Ci-CT-expressing cells around the stomach of *Ciona intestinalis*.** Consistent with a previous report[50], we observed cells (*) that express Ci-CT around the stomach of adult *C. intestinalis*. Expression detected by mRNA *in situ* hybridization by chain reaction on a transverse section through adult *C. intestinalis*. *st*, stomach.

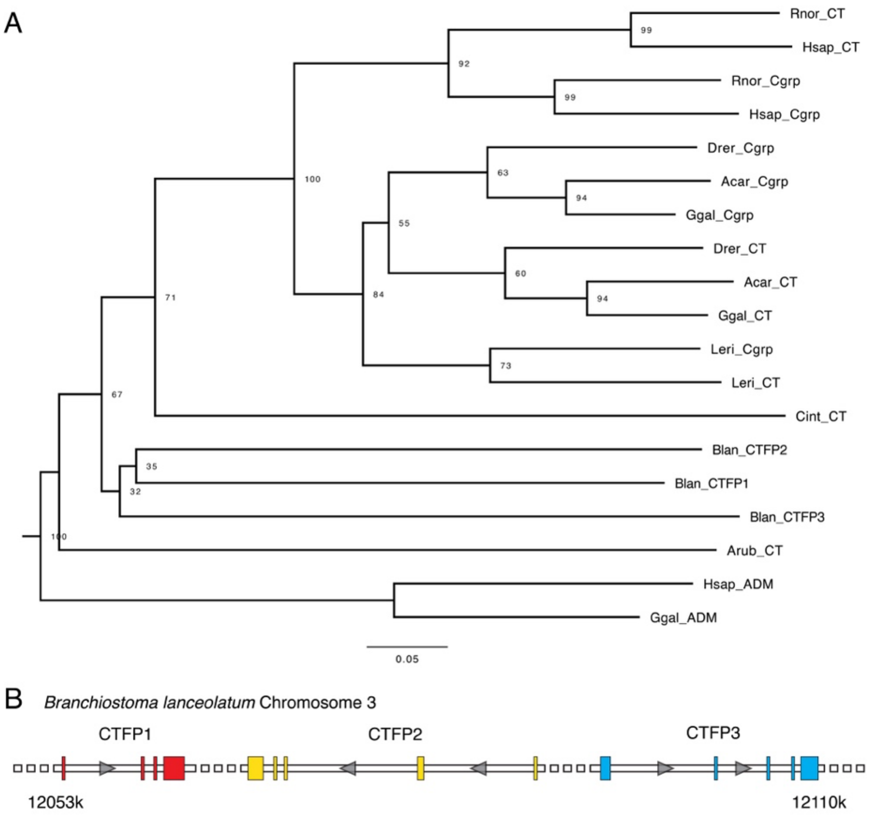


**Figure S4: Phylogeny of chordate calcitonin and calcitonin gene-related peptide sequences. a)** Neighbour joining molecular phylogeny of chordate calcitonin (CT) and calcitonin gene-related peptide (Cgrp) sequences reveals a lineage-specific triplication of the locus encoding the calcitonin family protein (CTFP) in the amphioxus, *Branchiostoma lanceolatum*. Sequences used for this phylogenetic analysis are listed in Table S1. **b)** Genes encoding *B. lanceolatum* CTFP1-3 are arranged in a cluster on chromosome 3. *Acar*, *Anolis carolinensis*; *Arub*, *Asterias rubens*; *Blan*, *Branchiostoma lanceolatum*; *Cint*, *Ciona intestinalis*; *Drer*, *Danio rerio*; *Ggal*, *Gallus gallus*; *Hsap*, *Homo sapien*; *Leri*, *Leucoraja erinacea*; *Rnor*, *Rattus norvegicus*.

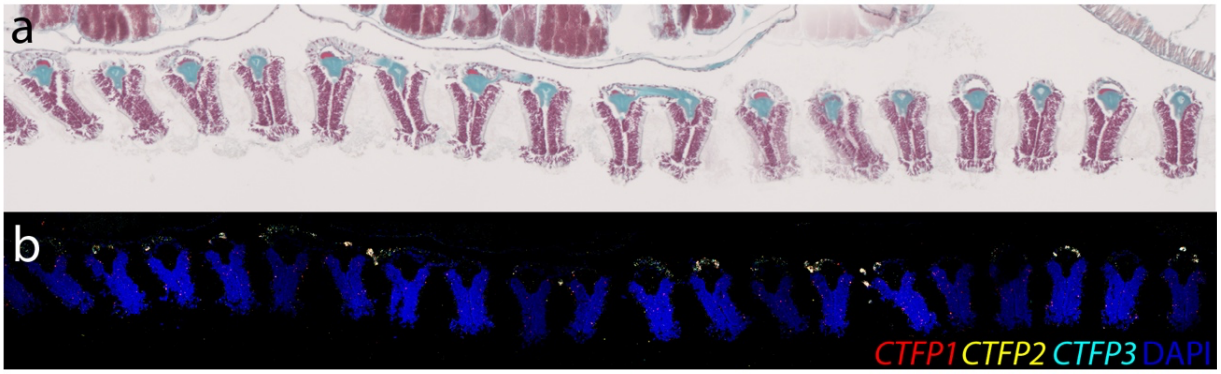


**Figure S5: CTFP expression in the pharyngeal arches of the amphioxus, *Branchiostoma lanceolatum*. a)** Low-power view of the pharyngeal bars of an adult *Branchiostoma lanceolatum*, stained with Masson's Trichrome. **b)** *CTFP1-3* are co-expressed in the atrial cells of each *B. lanceolatum* pharyngeal bar.
